# The pore-forming protein APOL7C ruptures *Leishmania major*-containing parasitophorous vacuoles in dendritic cells to promote IL-12 production and the induction of T helper type 1 immunity

**DOI:** 10.64898/2025.12.24.696354

**Authors:** Gerone Gonzales, Eymi Cedeño, Liam Wilkinson, Song Huang, Cassandra M. Wood, Neil McKenna, Robin M. Yates, Matheus B. Carneiro, Nathan C. Peters, Johnathan Canton

**Author notes:** These authors contributed equally (Gerone Gonzales, Eymi Cedeño).

## Abstract

Classical dendritic cells (cDCs) are required for the efficient priming of T helper type 1 (Th1) immunity against *Leishmania major* infections in mice. They do so by producing large quantities of IL-12 in response to *L. major* infection. IL-12 in turn supports the generation of protective CD4^+^ Th1 cells. However, the mechanism(s) by which *L. major* instigates robust IL-12 production by cDCs has not been addressed. Here we show that expression of a pore-forming protein called APOL7C is induced in cDCs by *L. major* infection. APOL7C is expressed in the cytosol of host cells and, remarkably, upon infection is recruited to *L. major* parasitophorous vacuoles (PVs) where it induces PV rupture. PV rupture leads to the exposure of *L. major*-derived pathogen-associated molecular patterns (PAMPs) to cytosolic pattern recognition receptors (PRRs). Cytosolic PRR signaling then drives IL-12 production by cDCs. Consequently, *L. major* challenge of APOL7C-deficient mice results in reduced numbers of IL-12 producing cDCs as well as a diminished Th1 response. In sum, our data indicate the presence of pore-forming proteins in the cytosol of cDCs that interfere with the intracellular niche of *Leishmania* to modulate immunity to these parasites.

## Introduction

Infection by protozoan parasites of the genus *Leishmania* results in Leishmaniasis – a disease affecting more than 12 million people across the world with 0.7 to 1 million new cases each year^1^. *Leishmania* parasites are deposited into mammalian hosts in their motile promastigote form by the bite of an infected sand fly. Once deposited, the promastigotes are engulfed by phagocytes such as neutrophils, monocytes, macrophages and classical dendritic cells (cDCs) through a process that closely resembles phagocytosis^2,3^. Immediately following engulfment, the parasites reside in membrane-bound compartments referred to as parasitophorous vacuoles (PVs). PVs are the primary site of parasite survival and replication in the mammalian host^2^. For this reason, a clear understanding of how host immune cells target the PV niche to control parasite survival and replication is of critical importance.

Shortly after they are formed, PVs, like phagosomes, undergo a series of interactions with endocytic organelles and lysosomes^4–6^. These interactions result in the acquisition of the NADPH oxidase that can generate antimicrobial reactive oxygen species (ROS), vacuolar(V)-ATPases that drive the acidification of the PV, hydrolases (including proteases, lipases and nucleases), and antimicrobial peptides^2,7^. The ability of *Leishmania* parasites to survive and indeed replicate within the seemingly hostile environment presented by the PV is determined by (i) the evasion mechanisms of the parasite and (ii) the activation status of the host cell^2,7^. The latter is dependent on the general inflammatory milieu at the site of infection and, later, is informed by signals from lymphocytes. For example, interferon γ (IFN-γ) from CD4+ T helper 1 (Th1) cells can stimulate phagocytes to effectively eradicate intracellular parasites via production of intraphagosomal reactive nitrogen species (RNS)^8–10^ . Indeed, a robust Th1 response is central to the control of *Leishmania* parasites *in vivo*. It is therefore not surprising that eliciting a more effective Th1 response has become the target of many therapeutic and vaccine strategies for leishmaniasis^11^.

Recent studies have demonstrated that *Batf3*-dependent cDCs (also known as Type 1 classical dendritic cells [cDC1s]) are essential players in the generation of robust Th1 responses to *Leishmania* parasites^12,13^. This was attributed to the release of large quantities of IL-12 by *Batf3*-dependent cDCs upon *Leishmania* infection^13^. Since this discovery however, the events leading to IL-12 production by *Batf3*-dependent cDCs in the context of *Leishmania* infection remain unclear. For example, IL-12 production during *Leishmania* infection is often assumed to result from the recognition of *Leishmania*-derived PAMPs by cell surface or endosomal TLRs^14,15^; but, there is growing evidence that *Leishmania*-derived PAMPs also trigger cytosolic PRRs, including some of which contribute to IL-12 production^16–20^. This presents a conundrum – how do *Leishmania*-derived PAMPs traverse intact PV membranes to gain access to cytosolic PRRs?

In this paper, we describe a previously unknown mechanism by which cDCs can target the intracellular niche of *Leishmania* parasites to elicit IL-12 production. Specifically, a cytosolic pore-forming protein called APOL7C is upregulated in cDCs during *Leishmania* infection and is recruited to PVs. Strikingly, arrival of APOL7C on PVs results in PV rupture. Disruption of PVs by APOL7C in turn leads to the exposure of *Leishmania*-derived PAMPs to cytosolic PRRs. Cytosolic PRR engagement leads to IL-12 production and supports a Th1 response. Our data therefore describes the existence of cytosolic pore forming effector proteins that are deployed by cDCs to mediate parasite control in the context of *Leishmania* infections.

## Results

### Classical dendritic cells (cDCs) at sites of *L. major* infection express the pore-forming protein APOL7C and are infected by *L. major*

Recently, we reported that cDCs, one of the cell types infected by *Leishmania* parasites *in vivo*, express a pore-forming protein called APOL7C^21^. Notably, we observed APOL7C recruitment to *L. major* PVs; however, we did not explore the consequence of APOL7C recruitment to PVs in cDCs. To address this, we first assessed whether APOL7C-expressing cells were present at sites of *L. major* infection. We re-analyzed a single cell RNAseq (scRNAseq) dataset from a previous study in which 2 X 10^6^ parasites were deposited intradermally into the ears of C57BL/6NCr mice and processed for scRNAseq after 4 weeks of infection (NCBI GEO; GSE185253)^22^. Seurat analysis revealed several CD45-positive clusters corresponding to monocytes (*Lst1, Ccr2, Ifitm3, Lyz2, Ly6c2*), macrophages (*Fcgr1, Csf1r, Mafb, Cybb, Adgre1*), T cells (*Cd3e, Cd28, Cd2, Cd3d, Cd40lg)*, neutrophils (*Csf3r, S100a8, S100a9, Mmp8, Mmp9*), cDCs (*Flt3, Dpp4, Zbtb46, Itgax, Ffar2*), CCR7^+^ cDCs (*Ccr7, Fscn1, Laptm4b, Socs2, Ncoa7*), mast cells (*Gata2, Cpa3, Ms4a2, Il1rl1, Fcer1a*), and proliferating cells (*Mki67, Stmn1, Top2a, Cdk1*) (**Fig 1A-D**)^21,23,24^. CD45-negative cells were not annotated as they did not express *Apol7c.* In the naïve mice, *Apol7c* expression was largely restricted to CCR7^+^ cDCs (**Fig 1E-F**). In infected mice, *Apol7c* expression was also present in CCR7^+^ cDCs and, notably, a small fraction of monocytes also appeared to express *Apol7c* (**Fig 1E-F**). However, compared to the entire population of monocytes, the proportion expressing *Apol7c* was relatively small (**Fig 1F**). Given the presence of *Apol7c*-expressing cDCs at sites of *L. major* infection, we next checked whether we could detect infection of cDCs in the ear during an *L. major* infection. For this, we intradermally inoculated 10^4^ *L. major* parasites that express RFP (*L. major*-RFP) into the ears of C57BL6/J mice and monitored cDC infection by flow cytometry. Using a previously validated flow cytometry panel^25,26^, we could readily detect infection of various intradermal DC subsets, including cDC1s (CD11B^-^, CD11C^+^, MHCII^+^, XCR1^+^), monocyte-derived dendritic cells (CD11B^+^, LY6G^-^, LY6C^-/Lo-Hi^, CD64^+^, CCR2^+^, MHCII^+^), a mixed population of heterogenous resident cDC2s and monocyte-derived DCs (henceforth referred to as Het-DCs; LY6C^-^, LY6G^-^, CCR2^+/-^, CD64^-^, CD11C^+^, MHCII^+^), and their CCR7^+^ equivalents, at the site of infection at 7 weeks post infection (**Fig 1G-H**, for gating strategy see **S1 Fig A-B**). After 18 weeks, the number of infected DCs was largely diminished (**Fig 1I**). Altogether our data demonstrate that various types of intradermal DCs at sites of *L. major* infection, including cDCs and CCR7^+^ cDCs, express *Apol7c* and that these *Apol7c*-expressing cells may be infected by *L. major*.

**Fig 1.**
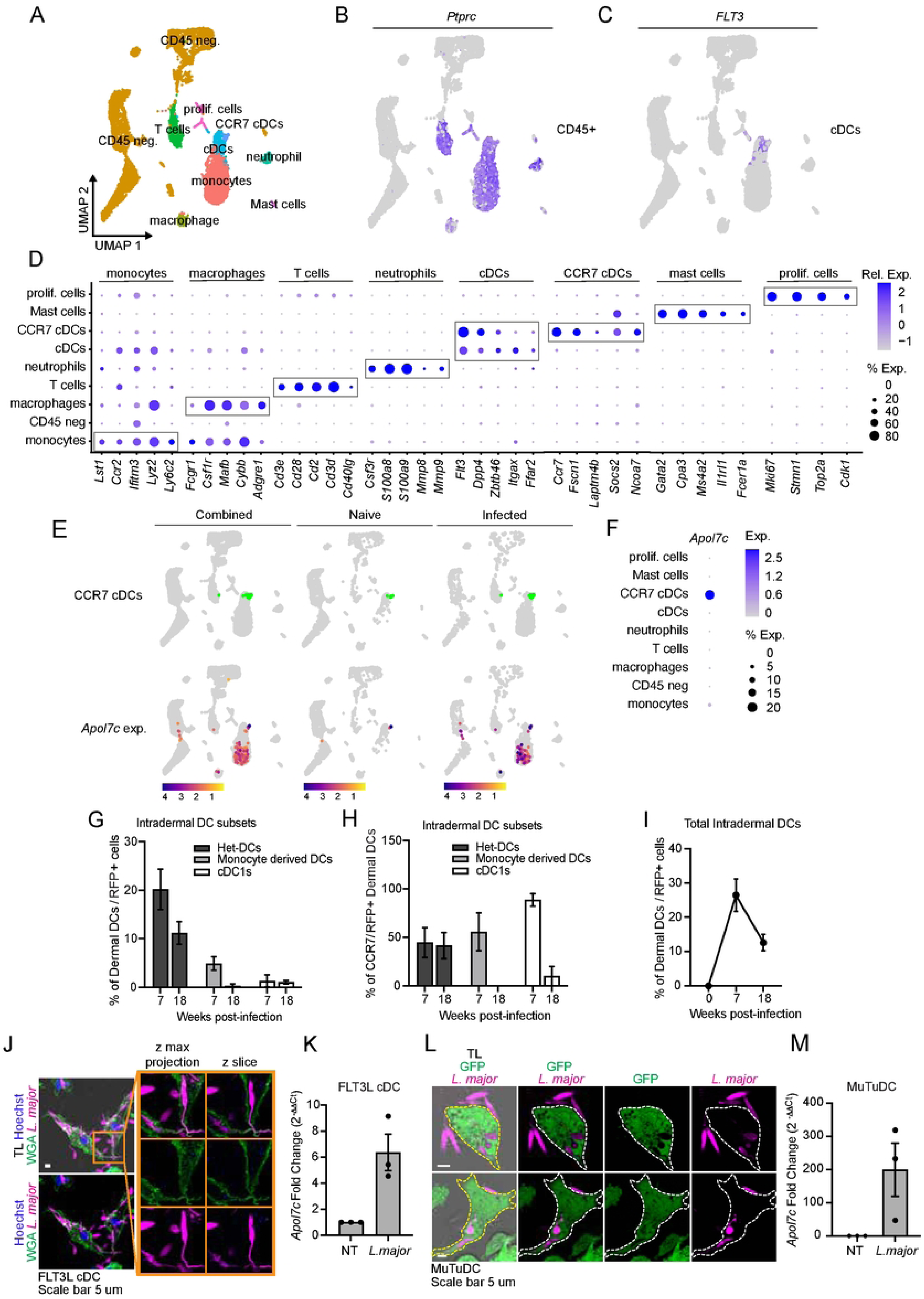
Infected cDCs at sites of *L. major* infection express *Apol7c*. *Apol7c* expression is enhanced in FLT3L cDCs and MuTuDCs upon *L. major* infection. **(A-C)** Uniform manifold approximation and projection (UMAP) of re-analyzed scRNAseq datasets of intradermally *L. major* infected ears showing **(A)** cell clusters of **(B)** CD45+ cells (expression of *Ptprc*) and **(C)** cDCs (expression of *FLT3*). **(D)** Curated genes used for annotations of cell clusters in **(A-C).** Colour indicates average expression of each gene (grey; low expression, blue; high expression). Sizes of circle represents the percent of cells within a cluster expressing the indicated genes. **(E)** UMAPs showing expression of CCR7+ cDCs and *Apol7c* expression pattern in naive and infected mice in cell clusters from **(A)**. **(F)** *Apol7c* expression in cell clusters from **(A)**. Colour indicates average expression of each gene (grey; low expression, blue; high expression). Sizes of circle represents the percent of cells within a cluster expressing the indicated genes. **(G-I)** C57BL/6J mice were intradermally inoculated with 10^4^ RFP-*L. major* parasites. At the indicated time points, flow cytometry was performed on single cell suspensions investigating DC populations at the site of infection. Data is plotted as mean ± SEM of 2 independent experiments (n=3-4). FLT3L cDCs **(J)** or MuTuDCs **(L)** were challenged with RFP expressing *L. major* at an MOI of 25. After 6 hours, cells were fixed, stained and imaged with confocal microscopy. FLT3L cDCs **(K)** or MuTuDCs **(M)** left untreated (NT) or challenged with *L. major* overnight were assessed for *Apol7c* expression by RT-qPCR. Plotted as mean ± SEM of 3 independent experiments (n=3). TL; transmitted light, WGA; wheat germ agglutinin-AF488.

### Exposure to *L. major* increases the expression *of Apol7c* by cDCs

We next sought to explore in greater detail the effect of *L. major* exposure on *Apol7c* expression by cDCs. We have previously found that innate immune stimulation of cDCs potently increases the expression of *Apol7c*^21^. To investigate whether the exposure of *L. major* to cDCs had any effect on *Apol7c* expression, we generated primary cDCs by culturing bone marrow cells with FLT3L and GM-CSF (henceforth FLT3L cDCs). FLT3L cDCs consist of a mixture of primary cDC1s and cDC2s and have previously been used to study APOL7C function^21,27^. FLT3L cDCs were challenged with *L. major*-RFP at an MOI of 25. Confocal imaging of cells stained with Wheat Germ Agglutinin (WGA)-AF488 to visualize the cell membrane revealed that FLT3L cDCs were readily infected by *L. major* (**Fig 1J**). Next, *Apol7c* expression was assessed by RT-qPCR. *L. major* infection resulted in strong upregulation of *Apol7c* by the FLT3L cDCs (**Fig 1K**). We also assessed the effect of *L. major* infection on *Apol7c* expression in the cDC1 cell line MuTuDC1940 (henceforth MuTuDCs). We have previously confirmed that MuTuDCs normally express *Apol7c* and respond to innate immune stimuli with an increase in *Apol7c* expression^21^. Like FLT3L cDCs, exposure of MuTuDCs to *L. major* also resulted in the strong upregulation of *Apol7c* (**Fig 1L-M**).

### APOL7C is recruited to LAMP1^+^ *L. major*-containing PVs

As mentioned above, we have shown that APOL7C is recruited to phagosomes containing diverse cargoes as well as to *L. major* PVs^21^. However, we did not investigate the kinetics and mechanism of APOL7C recruitment to PVs. To explore this in greater detail, we employed a previously established Raw264.7 cell line that can be induced to express APOL7C::mCherry or APOL7C::GFP upon treatment with doxycycline (henceforth RawKb.APOL7C::mCherry or RawKb.APOL7C::GFP)^21^. While this is not a cDC cell line, we have previously established that this cell line is suitable for the study of APOL7C recruitment to phagosomes as (i) we have shown that APOL7C::mCherry and APOL7C::GFP are recruited to phagosomes in these cells by the same mechanism as in primary cDCs^21^, (ii) Raw264.7 cells, unlike MuTuDCs and primary cDCs, lack endogenous *Apol7c* expression and therefore RawKb.APOL7C::mCherry cells represent a true gain-of-function system in which all APOL7C proteins are tagged with a fluorophore and the presence of APOL7C is strictly under the control of doxycycline^21^, (iii) Raw264.7 cells are amenable to genetic manipulation allowing for the expression of APOL7C mutants^21^ and (iv) Raw264.7 cells represent a widely accepted model for studying the cell biology of *Leishmania* PVs^28,29^. In uninfected cells, APOL7C::mCherry remained largely cytosolic; however, upon infection with *L. major,* APOL7C::mCherry redistributed to PV membranes after approximately 6 hours of infection (**Fig 2A-B, S1 Movie**). This late recruitment of APOL7C to PVs implied that APOL7C arrived on PVs that had already fused with lysosomes. We confirmed this by staining for LAMP1, a protein that is only present on lysosome membranes or on organelles that have fused with lysosomes. Indeed, APOL7C::GFP was only present on PVs that were also LAMP1^+^ (**Fig 2C-D**). Similarly, pre-loading lysosomes with fluorescent 10KDa dextran followed by *L. major* challenge revealed that greater than 75% of the APOL7C::mCherry-positive PVs were PVs that were dextran-positive and therefore had already fused with lysosomes (**Fig 2E-F**). Altogether these results indicate that APOL7C is recruited to PVs after they have fused with lysosomes.

**Fig 2.**
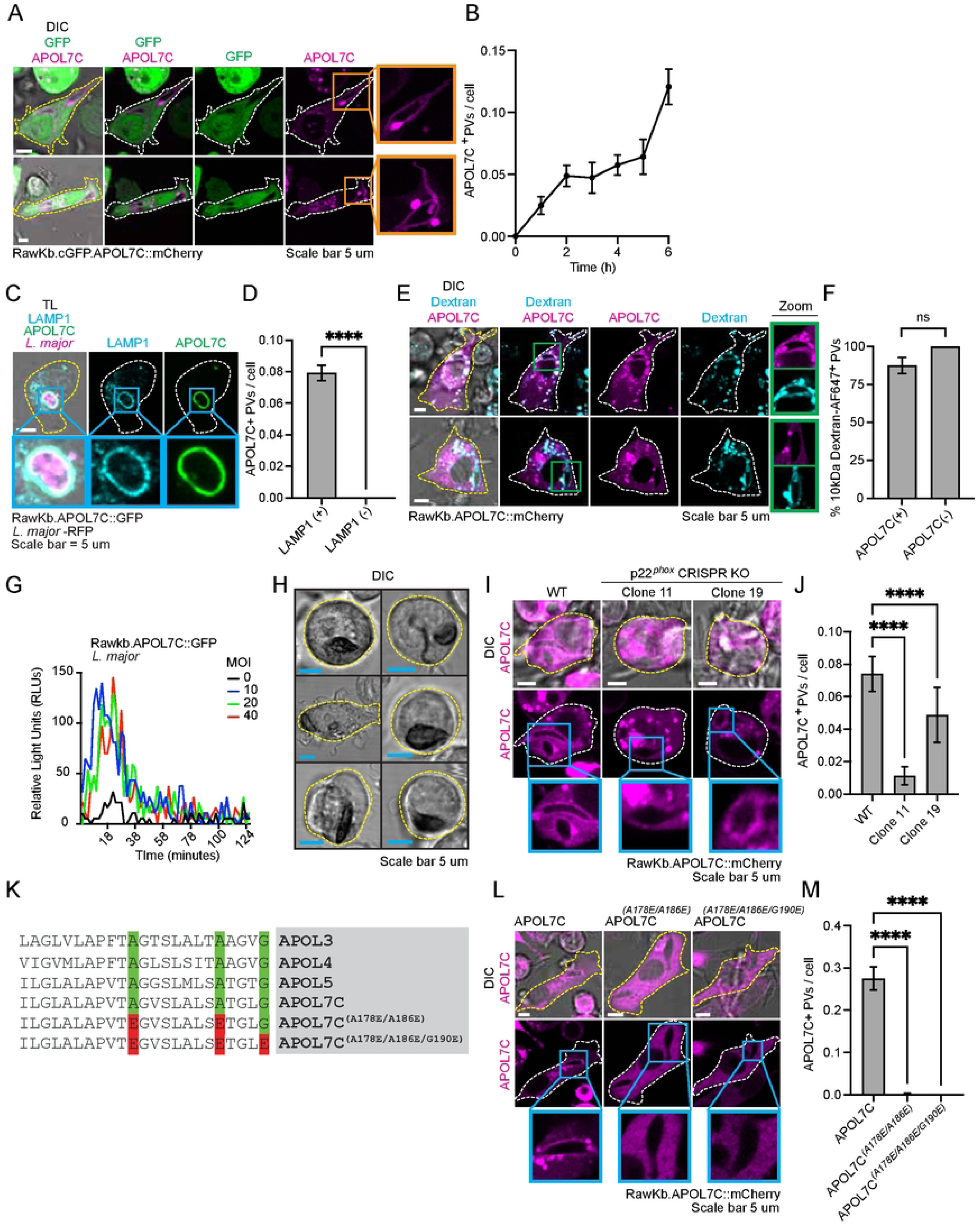
APOL7C is recruited to PV membranes fused with lysosomes and is NADPH oxidase- and voltage-dependent in gain-of-function experiments in Raw264.7 cells. (A-B) RawKb.cGFP.APOL7C::mCherry cells were plated and treated with doxycycline (1 μg/mL) overnight. The next day, cells were challenged with *L. major* with an MOI of 25, washed and imaged live with confocal microscopy at the indicated time points. **(B)** The number of APOL7C-positive PVs per cell were quantified. Plotted as mean ± SEM of three independent experiments (n=3). **(C-D)** RawKb.APOL7C::GFP cells were plated and treated with doxycycline overnight. The next day, cells were challenged with RFP-*L. major* with an MOI of 50 for hours before fixation and staining with LAMP1. Samples with then imaged with confocal microscopy and **(D)** the number of APOL7C-positive RFP-*L. major* harbouring PVs were quantified on LAMP1-positive or negative PVs. Plotted as mean ± SEM of three independent experiments (n=3), significance determined using student’s *t* test. **(E-F)** RawKb.APOL7C::mCherry cells were plated and treated with pulsed with 10 kDa dextran-AF647 for 6 hours. Fluorescent dextran was chased to lysosomes overnight along with treatment with doxycycline. The next day, cells were challenged with *L. major* with an MOI of 25, washed and imaged live with confocal microscopy after 3 hours. **(F)** the number of Dextran-positive PVs were quantified. Plotted as mean ± SEM of three independent experiments (n=3), significance determined using student’s *t* test. **(G)** RawKb.APOL7C::GFP cells were challenged with *L. major* parasites at the indicated MOIs in the presence of 50 μM luminol and 8 U/mL HRP. Chemiluminescence was measured every 2 minutes for a total of 120 minutes. Data is plotted as mean of experimental triplicates and is representative of 3 independent experiments (*n* = 3). **(H)** RawKb.APOL7C::mCherry cells were challenged with L. major parasites at an MOI of 25 for 4 hours. Extracellular parasites were then washed and cells were incubated in 1 μg/mL NBT for 15 minutes before live cell imaging. Dark formazan deposits indicate the production of ROS. **(I-J)** Wild-type (WT) and p22^phox^ KO RawKb.APOL7C::mCherry cells were plated and treated with doxycycline overnight. Cells were then challenged with *L. major* parasites for 4 hours, washed and imaged live. Data is plotted as mean ± SEM of three independent experiments (n=3), significance determined by a Kruskal-Wallis test comparing p22^phox^ KO and wild-type cells. **(K)** Alignment of APOL7C (residues 168 to 190) with APOL3-5. Green highlighted residues signify wild-type residues characteristic of voltage-sensitive recruitment of APOLs to membranes. Red highlighted residues signify mutated residues in APOL7C^(A178E/A186E)^ and APOL7C^(A178E/A186E/G190E)^ voltage-insensitive mutants. **(L-M)** RawKb.APOL7C::mCherry, RawKb.APOL7C^(A178E/A186E)^::mCherry, or RawKb.APOL7C^(A178E/A186E/G190E)^::mCherry cells were plated and treated with doxycycline overnight. The next day, cells were challenged with *L. major* parasites for 4 hours, washed and imaged live. Data is plotted as mean ± SEM of three independent experiments (n=3), significance determined by using a one-way ANOVA with a Kruskal-Wallis test comparing RawKb.APOL7C::mCherry and RawKb.APOL7C^(A178E/A186E)^::mCherry, or RawKb.APOL7C^(A178E/A186E/G190E)^::mCherry cells. TL = Transmitted light; DIC = differential interference contrast; ns = not significant; \*\*\*\**P* £ 0.0001.

### APOL7C is recruited to *L. major*-containing PVs in a voltage-dependent manner

We next explored the mechanism by which APOL7C::mCherry is recruited to phagosomes. Recruitment of APOL7C to phagosomes requires the triggering of the NADPH oxidase which leads to depolarization of the phagosome membrane allowing for the voltage-dependent insertion of APOL7C into the phagosome membrane^21^. We investigated whether a similar mechanism applied to its recruitment and insertion into PV membranes. First, we assessed whether *L. major* infection triggers detectable NADPH oxidase activity in RawKb.APOL7C::mCherry cells. Indeed, *L. major* infection triggered a robust oxidative burst at various MOIs as determined by luminol-based ROS-detection and formazan deposition in the presence of nitroblue tetrazolium (NBT) (**Fig 2G-H**). Next, we tested whether APOL7C::mCherry is recruited to phagosomes in cells that lack a functional NADPH oxidase. For this, we employed two separate clonal lines of RawKb.APOL7C::mCherry cells in which the p22*^phox^* subunit of the NADPH oxidase has been deleted. We have previously shown that these lines are incapable of producing an NADPH oxidase-driven oxidative burst^21^. In these cells, we observed diminished recruitment of APOL7C::mCherry to PVs suggesting that, akin to phagosomes containing inert particles^21^, a functional NADPH oxidase supports the recruitment of APOL7C::mCherry to PVs (**Fig 2I-J**). The voltage-dependence of APOL7C insertion into phagosome membranes maps to three neutral amino acid residues in a region of the protein called the membrane insertion domain (MID) (**Fig 2K**)^30–32^. Mutation of either 2 or 3 of these key amino acids into acidic residues results in the inability of APOL7C to insert into the membrane of phagosomes^21^. Using stable Raw264.7 cell lines that express APOL7C harboring mutations in either 2 (RawKb.APOL7C^A178E/A186E^::mCherry) or 3 (RawKb.APOL7C^A178E/A186E/G190E^::mCherry) of these key residues, we found that voltage-insensitive mutants could not be recruited to *L. major*-containing PVs (**Fig 2L-M**). Altogether our data demonstrate that APOL7C is recruited to *L. major*-containing PVs in an NADPH oxidase- and voltage-dependent manner.

### APOL7C recruitment to PVs leads to PV rupture

We next investigated the consequence of APOL7C recruitment to PVs. APOL7C contains a colicin-like pore forming domain that is capable of generating breaches in the delimiting membrane of phagosomes harboring inert particles^21^. We wondered whether the recruitment of APOL7C to PVs resulted in similar damage to the PV membrane. To this end, we infected RawKb.APOL7C::mCherry cells with *L. major* and assessed the incidence of ruptured PVs by staining for Galectin 3 – an established marker of damaged phagosomes and endocytic organelles^21,33–35^. Remarkably, APOL7C::mCherry recruitment to PVs appeared to coincide with Galectin 3 positivity (**Fig 3A-B**). The inverse was not true, as APOL7C::mCherry negative phagosomes were not Galectin 3 positive. Similarly, we pulsed the membrane impermeant dye Lucifer Yellow (LY) into the lumen of phagosomes by having it present in the medium during the infection. After washing away the extracellular LY, we were able to detect luminal LY in PVs. However, LY leaked out of APOL7C-positive PVs (**Fig 3C-D**). This is consistent with what we have previously observed in phagosomes harboring inert particles and is suggestive of PV membrane rupture instigated by APOL7C recruitment^21^.

**Fig 3.**
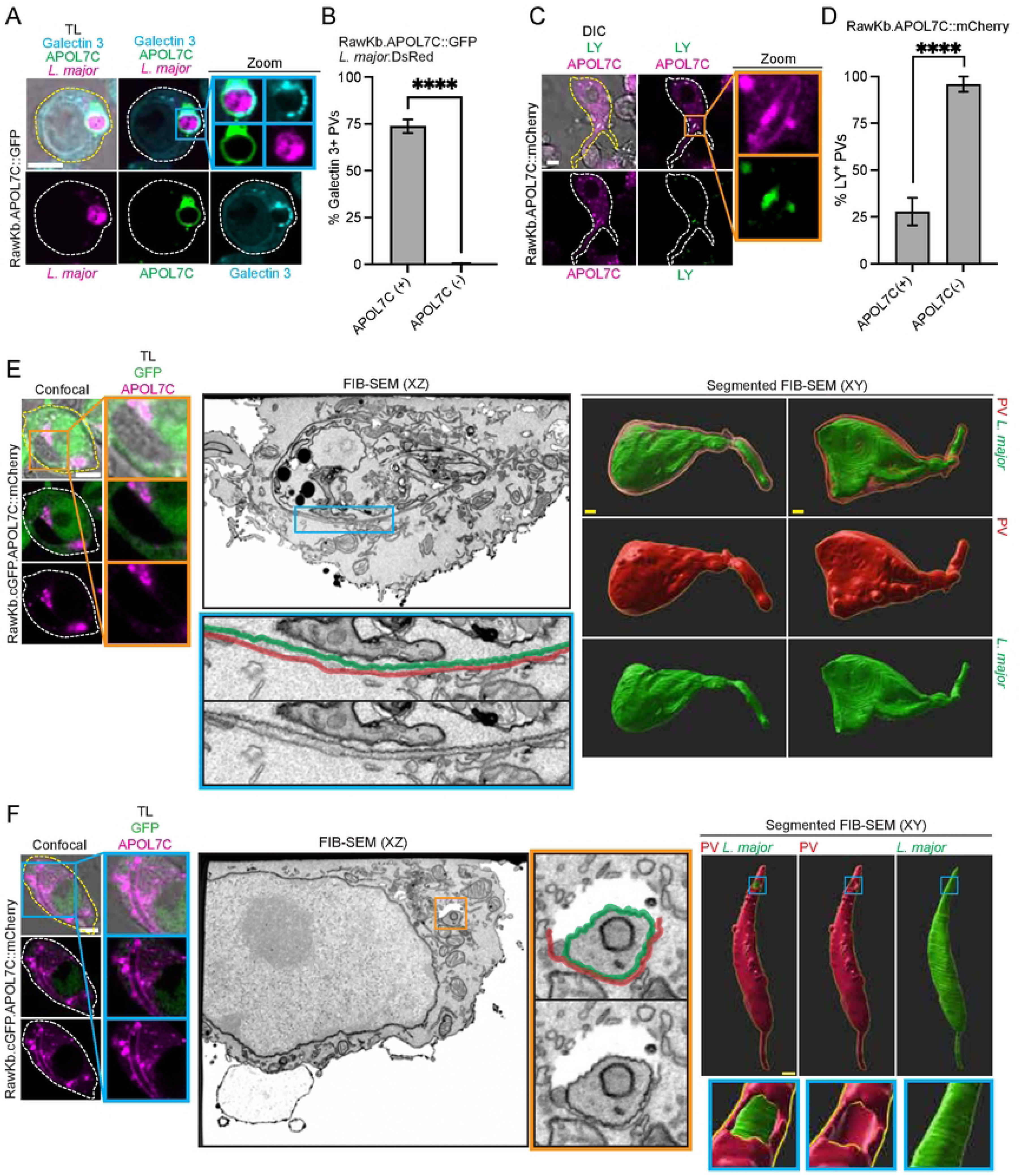
APOL7C on PVs induces membrane rupture. (A-B) RawKb.APOL7C::GFP cells were plated and treated with doxycycline (1 μg/mL) overnight. The next day, cells were challenged with RFP-*L. major* with an MOI of 50, fixed and stained for Galectin 3. Plotted as mean ± SEM of three independent experiments (n=3), significance determined using student’s *t* test. **(C-D)** RawKb.APOL7C::mCherry cells were plated and treated with doxycycline overnight. Cells were then challenged with *L. major* at an MOI of 25 with Lucifer Yellow (LY) in the media for 4 hours. The extracellular parasites and LY were then washed away and imaged with live confocal microscopy. The percentage of LY-positive PVs that were either APOL7C-positive or APOL7C-negative were quantified. Data is plotted as mean ± SEM of three independent experiments (n=3), significance determined using student’s *t* test. **(E-F)** RawKb.cGFP.APOL7C::mCherry cells were plated and treated with doxycycline overnight and challenged with *L. major* at an MOI of 25 for 4 hours. Cells with APOL7C-negative **(E)** or APOL7C-positive **(F)** PVs were then imaged live with confocal microscopy and processed for FIB-SEM. TL; Transmitted light, DIC; differential interference contrast; \*\*\*\**P* £ 0.0001.

To further investigate PV rupture by APOL7C during *L. major* infection, we performed correlative confocal microscopy and focused ion beam scanning electron microscopy (FIB-SEM). We observed discontinuous PV membranes indicative of membrane rupture in APOL7C-positive PVs (**Fig 3E, S2 Movie**), while APOL7C-negative PVs exhibited continuous PV membranes (**Fig 3F, S3 Movie**). Interestingly, we also observed damage to the parasite membrane in APOL7C-positive PVs (**S2 Fig**). Altogether our data show that the recruitment of APOL7C to PV membranes leads to the disruption of the parasite’s replicative niche.

### APOL7C-depedent PV rupture impairs intracellular survival of *L. major*

*L. major* has evolved to replicate within PVs^2,3,7,28,29^. We wondered whether APOL7C-dependent rupture may change the properties of the PV such that it becomes less hospitable to *L. major* survival and replication. We first assessed whether APOL7C-dependent PV rupture influenced intracellular parasite numbers. FLT3L cDCs from wildtype and *Apol7c^-/-^* mice were challenged with RFP-expressing *L. major* at an MOI of 50. Confocal microscopy of individual FLT3L cDCs revealed that *Apol7c^-/-^* cells had a larger parasite burden (**S3 Fig A-B**). However, the percentage of infected cells amongst bulk FLT3L cDCs remained unaffected by *Apol7c*-deficiency (**S3 Fig C**). We have previously shown that, in FLT3L cDCs, the primary cells expressing *Apol7c* are XCR1^+^ cDC1s^21^. Indeed, XCR1^+^ cDC1s from *Apol7c^-/-^* deficient mice appeared to have a higher parasite burden (**S3 Fig D-E**). Altogether, this data suggests that recruitment of APOL7C to *L. major* PVs likely disrupts the replicative niche and limits parasite replication.

### APOL7C-dependent PV rupture does not impact cross-presentation by cDCs

We have previously identified a critical role for APOL7C in cross-presentation by cDCs^21^. We therefore assessed whether APOL7C had any effect on the cross-presentation of *L. major*-derived antigens. FLT3L cDCs from wildtype and *Apol7c^-/-^* mice were challenged with ovalbumin(OVA)-expressing *L. major* at an MOI of 25. After four hours of infection, the extracellular parasites were washed away and the infected FLT3L cDCs were co-cultured with OT-I cells overnight. The medium was then assayed for OT-I-derived IFN-γ as an indirect measure of cross-presentation efficiency. While there appeared to be a slight defect in the cross-presentation of *L. major*-derived OVA in the APOL7C-deficient FLT3L cDCs, the difference was not significant (**Fig 4A**).

**Fig 4.**
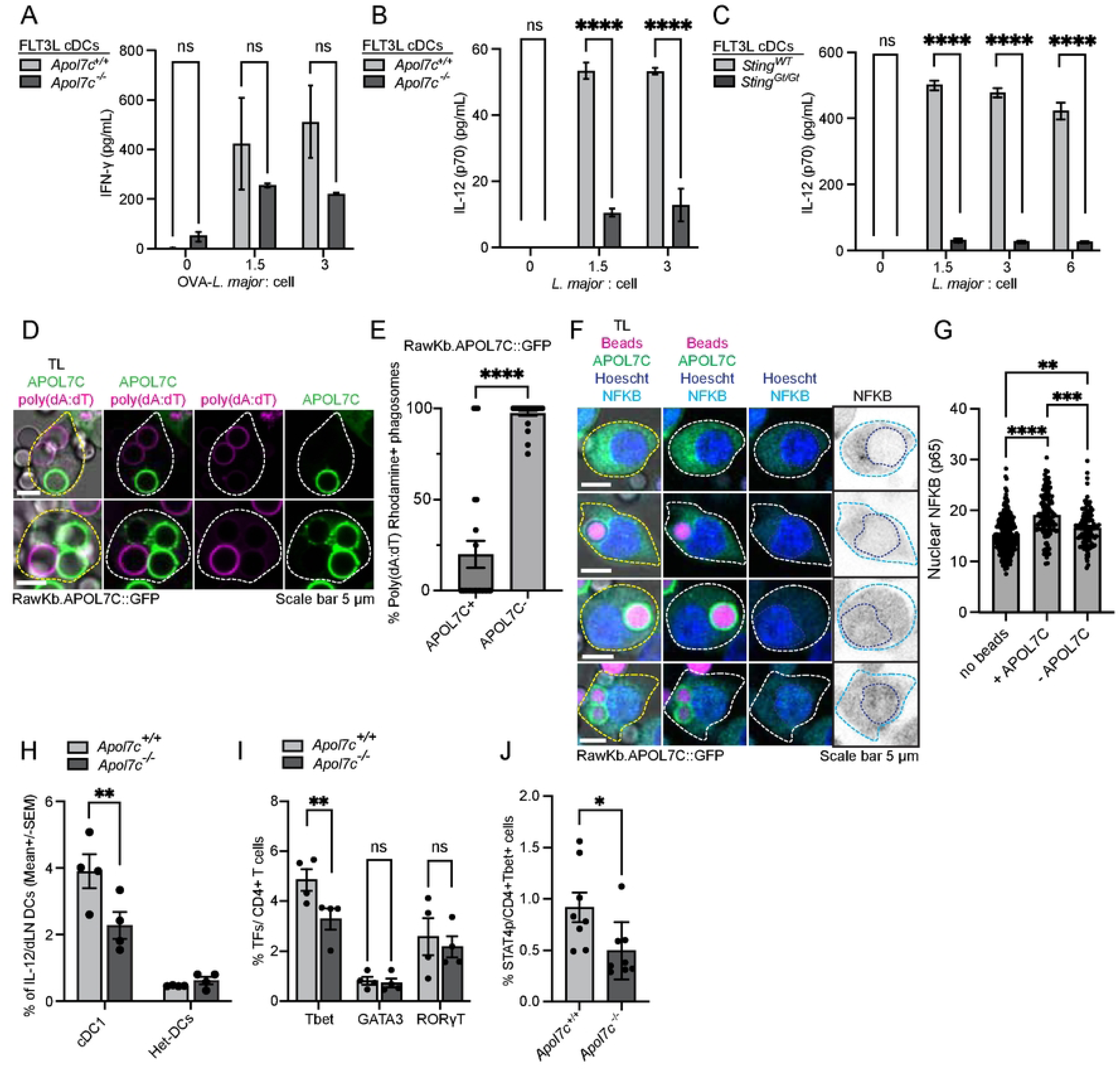
APOL7C mediated rupture promotes IL-12 production via leakage of PRR agonists. **(A)** FLT3L cDCs derived from *Apol7c^+/+^* and *Apol7c^-/-^* mice were plated and challenged with OVA-expressing *L. major* at the indicated MOIs overnight. Cells were then washed thoroughly and incubated overnight with OT-I cells. OT-I-derived IFN-g was measured by ELISA. Plotted as mean ± SEM of experimental triplicates and is representative of 3 independent experiments (*n* = 3). Significance determined using two-way ANOVA with Šídák’s multiple comparisons test. **(B-C**) FLT3L cDCs derived from **(B)** *Apol7c^+/+^* and *Apol7c^-/-^* mice or **(C)** *Sting^WT^* and *Sting^Gt/Gt^* mice were plated and challenged with *L. major* at the indicated MOIs overnight. Extracellular IL-12 was measured by ELISA. Plotted as mean ± SEM of experimental triplicates and is representative of 3 independent experiments (*n* = 3). Significance determined using two-way ANOVA with Šídák’s multiple comparisons test. **(D-E)** RawKb.APOL7C::GFP cells were plated and treated with doxycycline overnight. Cells were then challenged with non-porous silica beads and 50 μg/mL rhodamine-labelled poly(dA:dT) for 5 hours. Cells were then washed and imaged with live cell confocal microscopy. **(E)** The percentage of rhodamine-poly(dA:dT)-positive APOL7C-positive or APOL7C-negative phagosomes were quantified. Data is plotted as mean ± SEM of three independent experiments (n=3), significance determined using student’s *t* test. **(F-G)** RawKb.APOL7C::GFP cells were plated and treated with doxycycline overnight. Cells were then challenged with silica beads labelled with AF555 and 100 μg/mL naked poly(dA:dT) for 5 hours. **(G)** The nuclear stain was used to mask and quantify the mean fluorescence intensity (MFI) of NFKB in the nucleus of cells without internalized beads, and cells with APOL7C-positive and APOL7C-negative phagosomes. Data is plotted as mean ± SEM of three independent experiments (n=3), significance determined using a one-way ANOVA with a Kruskal-Wallis test. **(H-J)** *Apol7c^+/+^* and *Apol7c^-/-^* mice were intradermally inoculated with 2 X10^5^ RFP-*L. major* parasites for 48 hours. **(H)** IL-12-positive cDC1s and Het-DCs, **(K)** CD4+ T cells and **(L)** CD4+Tbet+ T cells from the draining lymph node (dLN) were then analyzed. **(H-I)** Data is plotted as mean ± SEM of data pooled from 4 mice per group (n=4) and significance was determined using two-way ANOVA with Šídák’s multiple comparisons test. **(J)** Data is plotted as mean ± SEM of data pooled from 8 mice (n=8) per group and Welch’s *t* test between *Apol7c^+/+^* and *Apol7c^-/-^* mice. TL = transmitted light; n.s. = no significance; \**P* £ 0.05; \*\**P* £ 0.01; \*\*\**P* £ 0.001; \*\*\*\**P* £ 0.0001.

### APOL7C-dependent PV rupture drives IL-12 production via cytosolic PRR signalling

While the finding that APOL7C deficiency had no impact on the cross-presentation of *L. major*-derived antigens was somewhat surprising, others have similarly shown that BATF3-deficient mice, which lack cDC1s, have no defect in the cross-presentation of *L. major*-derived antigens^13^. Instead, they found that cDC1s produce large quantities of IL-12 in response to *L. major* infection which contributes to a robust Th1 response^13^. Given that APOL7C is expressed by XCR1^+^ cDC1s in FLT3L cDCs^21^, we assessed whether IL-12 production by FLT3L cDCs was impacted by APOL7C deficiency. For this, FLT3L cDCs from wildtype and *Apol7c^-/-^* mice were challenged with *L. major* at an MOI of 25 overnight. The medium was then assayed for IL-12 by ELISA. Strikingly, APOL7C-deficient mice had a profound defect in their ability to elicit IL-12 production in response to *L. major* (**Fig 4B**). Based on this finding, we hypothesized that APOL7C-dependent rupture of PVs may expose *L. major*-derived PAMPs to cytosolic PRRs thereby enhancing IL-12 production by cDCs. While the engagement of cytosolic PRRs by *Leishmania*-derived PAMPs remains poorly characterized, recent work has shown that the cGAS-STING pathway is triggered during the infection of macrophage-like cells by *L. major*^36^. Specifically, kinetoplast DNA (kDNA), which can account for up to 30% of the parasite’s DNA, gains access to the cytosol and is sensed by the cGAS-STING pathway^36^. The cGAS-STING pathway is known to drive robust IL-12 production by cDCs^37,38^. We therefore assessed whether *L. major*-dependent IL-12 production by FLT3L cDCs was impacted by STING-deficiency. We generated FLT3L cDCs from wildtype and STING-deficient *Goldenticket* (*Sting^Gt/Gt^*) mice^39^. Remarkably, FLT3L cDCs from STING-deficient mice produced significantly less IL-12 relative to wildtype FLT3L cDCs in response to *L. major* infection (**Fig 4C**). This is consistent with the notion that APOL7C-dependent PV rupture may release PAMPs, such as kDNA, from within PVs to the cytosol to engage cytosolic PRR pathways such as the cGAS-STING pathway. To further investigate this possibility, we investigated whether poly(deoxyadenylic-deoxythymidylic) acid [henceforth poly(dA:dT)], an agonist of several cytosolic PRRs including cGAS, preferentially escaped from APOL7C-positive phagosomes. To this end, fluorescently-tagged poly(dA:dT) was pulsed into phagosomes in RawKb.APOL7C::GFP-expressing cells. Confocal microscopy revealed that phagosomes where APOL7C::GFP was recruited failed to retain poly(dA:dT) (**Fig 4D-E**). This is consistent with our previous work showing that APOL7C recruitment to phagosomes results in permeabilization of the phagosomes and subsequent leakage of the phagosomal contents to the cytosol. Next, we assessed whether the APOL7C-induced leakage of poly(dA:dT) to the cytosol would trigger inflammatory signalling ultimately leading to IL-12 production. For this, we chose to monitor the translocation of the p65 subunit of NF-κB from the cytosol to the nucleus. The triggering of cytosolic PRRs, such as cGAS, leads to p65 translocation and, importantly, this is a known driver of IL-12 production. We reasoned that as poly(dA:dT) leaks out of APOL7C-positive phagosomes it will engage cytosolic PRRs leading to p65 translocation to the nucleus. First, to ensure we could accurately measure p65 translocation, we measured nuclear translocation of p65 after stimulation of the RawKb.APOL7C::GFP with LPS – a TLR4 agonist and potent inducer of p65 translocation. As expected, we detected robust re-localization of p65 to the nucleus (**S4 Fig A-B**). Then, we loaded phagosomes with poly(dA:dT), as above, and assessed the nuclear translocation of p65 in cells with one or more APOL7C::GFP-positive phagosomes and in cells with no APOL7c::GFP-positive phagosomes. Remarkably, cells with APOL7C::GFP-positive phagosomes displayed the strongest translocation of p65 (**Fig 4F-G**). This is consistent with our finding that APOL7C::GFP-positive phagosomes leak poly(dA:dT) to the cytosol where it can engage cytosolic PRRs. Taken together, these data suggest that APOL7C recruitment to PVs and phagosomes results in their rupture and the leakage of PV or phagosomal contents to the cytosol. This in turn triggers cytosolic PRRs thereby driving inflammatory signalling and IL-12 production.

### APOL7C-deficient mice have reduced IL-12-producing cDCs in their draining lymph nodes (dLNs) after infection with *L. major*

Finally, we assessed whether infection of APOL7C-deficient mice with *L. major* resulted in a similar reduction in IL-12-producing cDCs relative to wild-type mice. For this, we intradermally inoculated 2 X10^5^ *L. major* parasites that express RFP (*L. major*-RFP) into the ears of C57BL6/J mice and, after 48 hours, cells from the site of infection and the draining lymph nodes (dLNs) were harvested for flow cytometry analysis. While we did not detect noticeable differences in the number of parasites at both the site of infection and the dLNs (**S4 Fig C-D**), we did observe a significant reduction in the number of IL-12-producing cDC1s, but not IL-12-producing Het-DCs, in the dLNs of APOL7C-deficient mice (**Fig 4H**, gating strategy in **S4 Fig E**). We also observed a significant reduction in TBET^+^ CD4^+^ T (Th1) cells in the dLNs of APOL7C-deficient mice while GATA3^+^ CD4^+^ T (Th2) cells and RORγT^+^ CD4^+^ T (Th17) cells were unaffected by APOL7C deficiency (**Fig 4I**, gating strategy in **S4 Fig F-H**). Similarly, the number of TBET^+^ CD4^+^ T cells with phosphorylated STAT4 (pSTAT4) was also reduced in APOL7C-deficient mice (**Fig 4J**, gating strategy in **S4** **Fig 4F and H**). STAT4 is phosphorylated downstream of cytokines such as IL-12 and is required for efficient Th1 polarization. Altogether, our data suggest that APOL7C-dependent release of *Leishmania*-derived PAMPs to the cytosol enhances IL-12 production by cDCs both *in vitro* and *in vivo*. This in turn favors a Th1 response to the parasite infection.

## Discussion

Some eukaryotic parasites, including several species of *Leishmania*, *Toxoplasma*, and *Trypanosoma*, have evolved strategies to survive in membrane-bound, intracellular compartments^40^. These compartments often represent privileged sites of parasite replication. Furthermore, the parasites can subvert host cell processes to establish these replicative niches. In the case of *Leishmania*, PVs resemble phagolysosomes in that they are acidic, contain various hydrolases with acidic pH optima and fuse extensively with lysosomes^2,3,7,29^. They differ from phagolysosomes in that *Leishmania* parasites can (i) alter the production of ROS within the PV by inactivating the phagocyte oxidase^41–44^, (ii) impact the rate at which PVs fuse with lysosomes^45,46^, and (iii) mediate the delivery of material from other membranous organelles, such as the ER, to the PV^28,29,40^. It is generally believed that their sequestration within these compartments facilitates evasion of the host immune response by limiting host detection and effector mechanisms^2,47^. Nevertheless, *Leishmania* can be eliminated by host immune cells and the mechanisms by which they do so are an active area of interest. In this manuscript, we have described previously unrecognized cytosolic effector proteins, called APOLs, with which phagocytes may target the replicative niche of intracellular parasites to favor parasite detection by the host and to modulate immunity.

The APOLs represent an understudied family of proteins that appear to play roles in immunity^21,48–51^. The family can be roughly divided into two groups: (i) secreted APOLs which contain a signal peptide and operate distally from host cells and (ii) cytosolic APOLs that remain within host cells. Much less is known about the latter group. However, we have recently described a role for the cytosolic APOL - APOL7C^21^. APOL7C is strongly induced by innate immune stimuli and is recruited to phagosomes where it changes the properties of the phagosome membrane. Ultimately, APOL7C recruitment to phagosomes leads to their rupture and results in the release of phagosomal contents to the cytosol. Here we found that in addition to phagosomes harboring inert particles, APOL7C can be recruited to *Leishmania*-containing PVs.

As with phagosomes, APOL7C instigates PV rupture leading to the escape of PV contents to the cytosol. Our previous work has shown that APOL7C-dependent rupture of phagosomes can lead to the escape of ingested antigens from within phagosomes to the cytosol for the efficient cross-presentation of exogenous antigen by dendritic cells^21^. However, we did not find any impact of APOL7C-dependent PV rupture on the cross-presentation of *Leishmania*-derived antigens. We cannot formally rule out an effect on cross-presentation because PV rupture appears to have a direct impact on the intracellular survival and might also affect the development of the parasites. It is likely that this impacts the availability of *Leishmania*-derived antigen making comparisons of cross-presentation by wildtype and APOL7C-deficient cDCs difficult to interpret.

Somewhat unexpectedly, we found that APOL7C-dependent rupture promotes the production of IL-12 by infected cDCs. This is consistent with previous work demonstrating the importance of cDC-derived IL-12 in the production of a robust and protective Th1 response to *L. major* infection^13^. We propose that APOL7C-dependent rupture leads to the exposure of *Leishmania*-derived PAMPs to the cytosol to elicit IL-12 production via cytosolic PRR signalling pathways. In support of this, we have presented evidence for the cGAS-STING pathway in the generation of IL-12 downstream of APOL7C-dependent PV rupture. However, we emphasize that there are other pathways leading to IL-12 production during *Leishmania* infection that likely do not require APOL7C-dependent PV rupture^14^. Indeed, APOL7C-deficient mice do still produce IL-12, albeit greatly reduced. Similarly, cGAS is not likely to be the only cytosolic PRR engaged following PV rupture. Indeed, our data demonstrates that in cDCs deficient in STING, IL-12 production is not completely abolished. Therefore, while APOL7C-dependent PV rupture strongly enhances IL-12 production by cDCs, it is but one pathway leading to IL-12 production by cDCs in the context of *L. major* infection. This may be the reason that, despite reduced IL-12 production by cDCs *in vivo* and reduced numbers of Th1 cells, we did not detect any reduction in parasite numbers. The IL-12 produced by alternate APOL7C-independent pathways may have been sufficient to control parasite numbers. The redundancy in host pathogen detection pathways that lead to IL-12 production during *Leishmania* infection likely evolved to ensure parasite clearance despite parasite-dependent evasion of individual host detection pathways. Finally, it is worth noting that there is a very large recruitment of monocytes to the *L. major* infection site, it is possible that IL-12-producing monocytes may take the place of APOL7C-expressing cDCs as the major source of IL-12 as the infection progresses^52^. Nevertheless, the evidence we present demonstrates an important role for APOL7C-dependent PV rupture for IL-12 production by cDCs early on in the infection.

This study focused on APOL7C which is expressed by dendritic cells and to a lesser extent by some monocytes^21^. The primary host cells of *Leishmania* parasites are believed to be monocytes, monocyte-derived macrophages and tissue-resident macrophages. While the latter are not the primary expressors of APOL7C, it is worth noting that other members of the APOL family are preferentially expressed in monocytes and macrophages. Notably, several publicly available datasets, including Gainullina *et al.*^53^, demonstrate selective expression of APOL9A and APOL9B in Kupffer cells^53^. Kupffer cells are a major reservoir of replicating *Leishmania* parasites in the case of visceral leishmaniasis^54^. It is conceivable that the expression of other APOLs, such as APOL9A and APOL9B, in other, non-cDC host cells, may play a similar role to APOL7C in IL-12 production in the context of *Leishmania* infection.

Altogether, we have identified pore-forming effectors of innate immune cells that allow for the disruption of *L. major*-containing PVs. Akin to guanylate-binding proteins (GBPs)^55^, APOLs may serve as a second line of defense for intracellular parasites that are resistant to intra-phagosomal killing. At the level of individual immune cells, PV rupture restricts parasite survival and replication; however, like GBPs^55^, PV rupture also unveils *L. major* to cytosolic innate immune sensors which in the *in vivo* setting leads to the modulation of adaptive immunity.

## Materials and Methods

### General materials used in this study

| Product | Catalogue number | Company |
| --- | --- | --- |
| RPMI1640 | SH30027.01 | Cytiva |
| IMDM | SH30259.02 | Cytiva |
| DMEM | SH30022.01 | Cytiva |
| DPBS | SH30028.03 | Cytiva |
| Fetal bovine serum (FBS) | 2027-06 | Corning |
| Bovine growth serum (BGS) | SH30541.03 | Cytiva |
| β-mercaptoethanol | 31350-010 | Gibco |
| Pen Strep solution | SH30010 | Cytiva |
| Puromycin | A11138-03 | Gibco |
| Hygromycin | Ant-hg-1 | Invivogen |
| Dimethyl sulfoxide (DMSO) | 0231-500ML | VWR |
| HEPES | 25060125 | Corning |
| M199 (1X) | 12350-039 | Gibco |
| M199 (10X) | 11825-015 | Gibco |
| Adenine | A8751 | Sigma |
| Hemin | 51280-5G | Fluka |
| Biotin | B5639-1G | Sigma |
| L-glutamine | 35050-061 | Gibco |
| Biopterin | B2517 | Sigma |
| Triethanolamine | 470302-964 | Ward's Science |
| Paraformaldehyde (16 %) | 15710 | Electron Microscopy Sciences (EMS) |
| Glutaraldehyde (8 %) | 16019 | Electron Microscopy Sciences (EMS) |
| In-Fusion HD Cloning Kit | 638948 | Takara |
| Prolong diamond anti-fade mountant | P36961 | Invitrogen |
| Alexa Fluor 555, SE | A20109 | Invitrogen |
| Silica beads, COOH modified | PSI-3.0COOH | Kisker Biotech |
| Silica microspheres | SS05003 | Bangs Laboratories, Inc |
| Fugene HD transfection reagent | 104162 | Active Motif |
| Zombie UV Live/Dead stain | 423107 | Biolegend |
| Zombie Red Live/Dead stain | 423109 | Biolegend |
| Zombie Green Live/Dead stain | 423111 | BioLegend |
| Coverslips (12 mm) | 48382 041 | VWR |
| Glass slides | 16004-388 | VWR |
| 8 well live cell imaging dish | 80826 | Ibidi |
| μ-dish gridded (35 mm) | 81168 | Ibidi |
| GlutaMAX supplement | 35050-061 | Gibco |
| Glycine | BP381-5 | Fisher |
| Cyanimide | 187364-25G | Sigma Aldrich |
| Ammonium chloride | A0171-500G | Sigma Aldrich |
| Lucifer yellow | L453 | Invitrogen |
| Doxycycline | J60579-14 | Alfa Aesar |
| Alt-R S.p. Cas9 Nuclease V3 | N/A | Integrated DNA Technologies (IDT) |
| Alt-R electroporation enhancer | N/A | Integrated DNA Technologies (IDT) |
| Sodium Bicarbonate solution | 17-613E | Lonza |
| GeneJet RNA purification kit | K0731 | ThermoFisher |
| EasySep Pan-DC enrichment kit II | 19863 | StemCell |
| qScript cDNA synthesis kit | 95047 | QuantaBio |
| iQ SYBR Green Supermix | 1708880 | BioRad |
| Osmium tetroxide, 4% solution | 19170 | Electron Microscopy Sciences |
| Potassium ferrocyanide | 470302-126 | Ward's Science |
| Durcupan ACM, Comp. A | 44611 | Sigma Aldrich |
| Durcupan, Comp. B | 44612 | Sigma Aldrich |
| Durcupan, Comp. C | 44613 | Sigma Aldrich |
| Durcupan, Comp. D | 44614 | Sigma Aldrich |
| Sodium Cacodylate Buffer, 0.2M | 11652 | Electron Microscopy Sciences |
| PELCO conductive silver paint | 16062 | Electron Microscopy Sciences |
| Carbenicillin | C1389 | Sigma Aldrich |
| Overnight express instant TB medium | 71491 | Sigma Aldrich |
| Protease inhibitor cocktail powder | P2714 | Sigma Aldrich |
| DNase I | 11284932001 | Sigma Aldrich |
| Lysoszyme | L6876 | Sigma Aldrich |
| Zwittergent 3-14 | T2653 | TCI America |
| His60 Ni superflow resin | 635659 | Takara |
| ELISA MAX Deluxe mouse IFN-γ kit | 430804 | BioLegend |
| ELISA MAX Deluxe mouse IL-12 (p70) kit | 433604 | BioLegend |
| 10 kDa dextran, AF647 | D22914 | ThermoFisher |
| Luminol | 521-31-3 | Sigma Aldrich |
| Nitro blue tetrazolium chloride | 0329-500MG | VWR |
| Poly(dA:dT) naked | tlrl-patn | InvivoGen |
| Poly(dA:dT) Rhodamine | tlrl-patrh | InvivoGen |
| Hoechst 33342 | 14533-100MG | Sigma Aldrich |

*Plasmids used in this study*.

| **Plasmid name** | **Source** | **Catalogue number** |
| --- | --- | --- |
| P40(PX)::GFP | Sergio Grinstein's lab | N/A |
| pSBtet-hyg | Addgene | 60508 |
| pSBtet-hyg-APOL7C::mCherry | Our lab, generated for this paper. | N/A |
| pcDNA3.1-APOL7C::mCherry | Our lab, generated for this paper. | N/A |
| pCMV(CAT)T7-SB100 | Addgene | 34879 |
| pSBbi-pur-H-2K<sup>b</sup> | Addgene | 111623 |
| pSBbi-GB | Addgene | 60520 |
| pSBtet-hyg-APOL7C<sup>A178E/A186E</sup>::mCherry | Our lab, generated for this paper. | N/A |
| pSBtet-hyg-APOL7C<sup>A178E/A186E/G190E</sup>::mCherry | Our lab, generated for this paper. | N/A |
| pSBtet-hyg-APOL7C::GFP | Our lab, generated for this paper. | N/A |

*Antibodies used in this study.* *Unconjugated primary antibodies.*
| Target | Clone | Stock | Application | Company |
| --- | --- | --- | --- | --- |
| LAMP1 | 1D4B | 0.5 mg/mL | IF 1:50 | Biolegend<br>121602 |
| mCherry | 8C5.5 | 0.5 mg/mL | IF 1:500 | Biolegend<br>677702 |
| p22<sup>phox</sup> | E9N8W | - | IF 1:100 | Cell Signaling<br>Technology<br>(CST) 88048S |
| NFKB (p65) | D14E12 | 139ug/mL | IF 1:400 | Cell Signaling<br>Technology<br>8242S |

*Antibodies used in this study. Unconjugated primary antibodies*.

*Conjugated primary antibodies*.

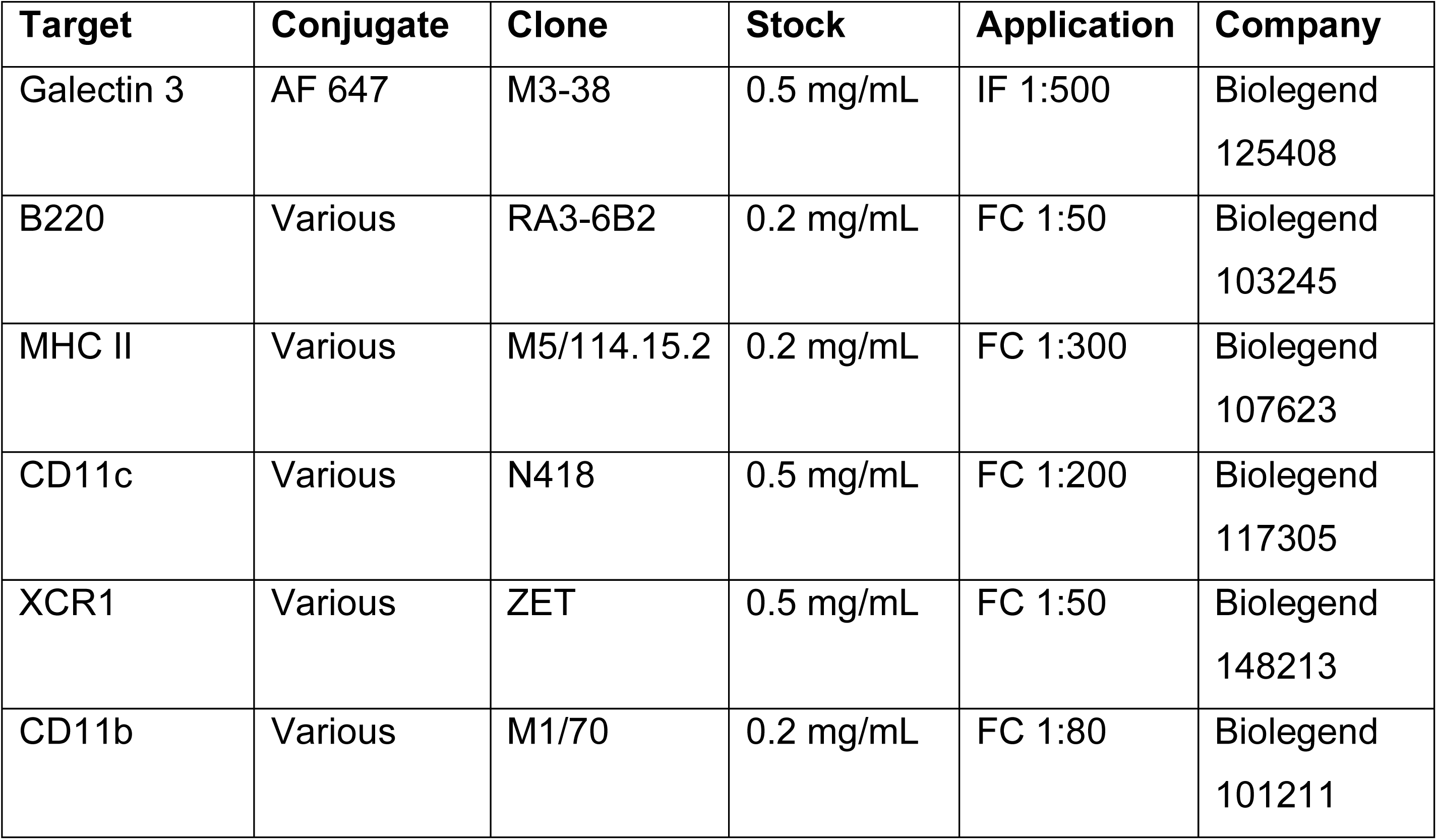

| Target | Conjugate | Clone | Stock | Application | Company |
| --- | --- | --- | --- | --- | --- |
| Galectin 3 | AF 647 | M3-38 | 0.5 mg/mL | IF 1:500 | Biolegend<br>125408 |
| B220 | Various | RA3-6B2 | 0.2 mg/mL | FC 1:50 | Biolegend<br>103245 |
| MHC II | Various | M5/114.15.2 | 0.2 mg/mL | FC 1:300 | Biolegend<br>107623 |
| CD11c | Various | N418 | 0.5 mg/mL | FC 1:200 | Biolegend<br>117305 |
| XCR1 | Various | ZET | 0.5 mg/mL | FC 1:50 | Biolegend<br>148213 |
| CD11b | Various | M1/70 | 0.2 mg/mL | FC 1:80 | Biolegend<br>101211 |

*Conjugated secondary antibodies*.

| Target | Conjugate | Species | Stock | Application | Company |
| --- | --- | --- | --- | --- | --- |
| Rabbit IgG | AF 488 | Goat | 2 mg/mL | IF 1:1000 | Invitrogen<br>A11034 |
| Rat IgG | AF 647 | Goat | 2 mg/mL | IF 1:1000 | Invitrogen<br>A21247 |
| Mouse IgG | AF 555 | Goat | 2 mg/mL | IF 1:1000 | Invitrogen<br>A21424 |

*Cells.* RPMI 1640 supplemented with 10% heat-inactivated bovine growth serum (HI-BGS), penicillin, and streptomycin was used to grow Raw264.7 and all derivatives of Raw264.7 cells. Raw264.7 cells were provided by Dr. Robin Yates (University of Calgary). The generation of the RawKb.APOL7C::mCherry, RawKb.APOL7C::GFP, RawKb.cGFP.APOL7C::mCherry, p22*^phox^*^-/-^ RawKb.APOL7C::mCherry, RawKb.APOL7C^A178E/A186E^::mCherry, and RawKb.APOL7C^A178E/A186E/G190E^::mCherry cell lines is described in Gonzales, Huang and Wilkinson *et al*. IMDM supplemented with 10% heat-inactivated fetal bovine serum (FBS), penicillin, streptomycin, glutaMAX, sodium bicarbonate and HEPES was used to grow MuTuDC1940 cells. MuTuDC1940 cells were purchased from Applied Biological Materials Inc. (ABM). Generation of the MuTuDC.APOL7C::mCherry cell line is described in Gonzales, Huang, Wilkinson *et al*. FLT3L-cDCs were generated as in Gonzales, Huang, Wilkinson *et al*. Specifically, C57BL/6 bone marrow derived cells were cultured in IMDM supplemented with 10% heat-inactivated fetal bovine serum (FBS), penicillin, streptomycin, glutaMAX, sodium bicarbonate, HEPES, 1.5 ng/mL GM-CSF and 60 ng/mL FLT3-L for 12-14 days. The culture medium was topped up at day 5 and cells were re-plated with fresh cytokines on day 9 and used between days 12-14.

*Mice.* C57BL/6J, OT-I, and *Batf3^-/-^* mice were obtained from Jackson. *Apol7c^-/-^* were generated by GemPharmatech and described in detail in Gonzales, Huang, Wilkinson *et al*. All mice were bred at the University of Calgary Foothills Campus mouse facility under specific pathogen-free conditions. Mice were used at 6-8 weeks of age and both male and female littermates were randomly assigned to experimental groups. Mice were born at normal Mendelian ratios. All mouse experiments were performed in accordance with national and institutional guidelines for animal care under specific protocols that underwent institutional review.

### Primers used to genotype mice

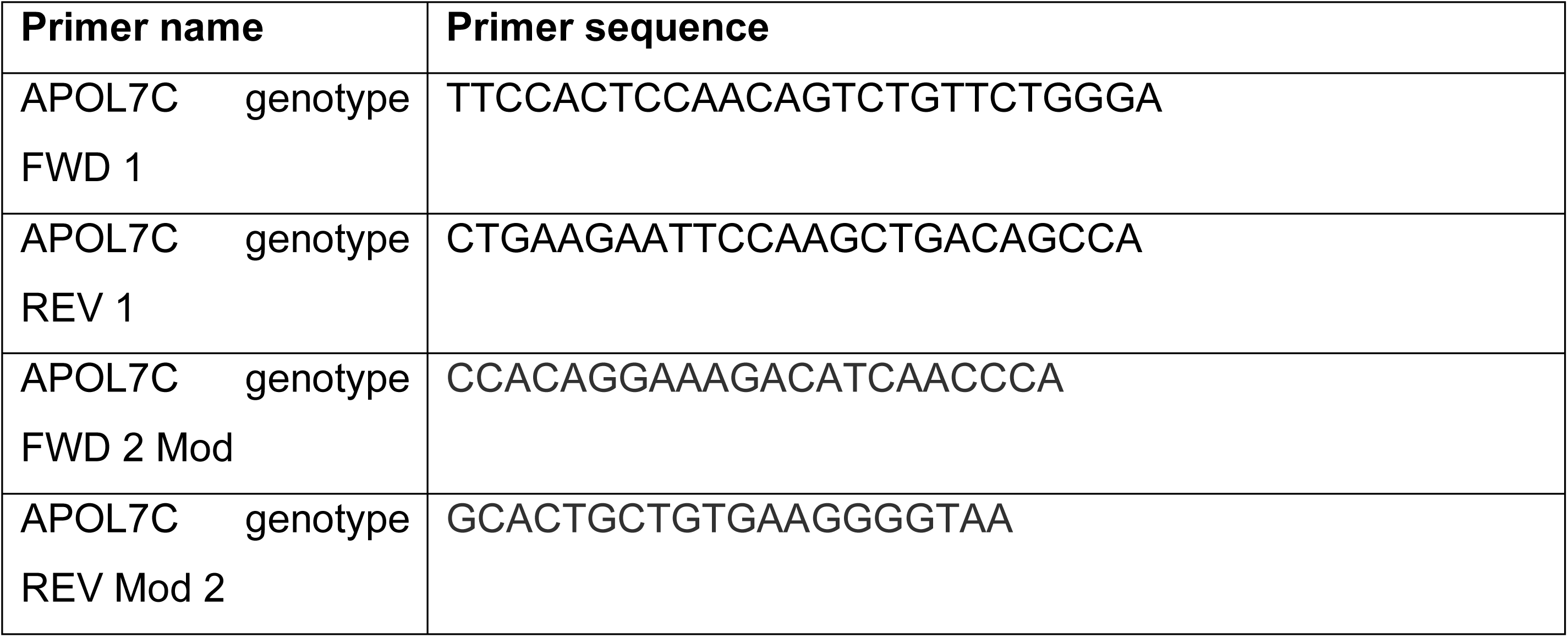

*RT-qPCR for APOL7C expression*. The mRNA was isolated using the Genejet RNA purification kit in accordance with the manufacturer’s instructions. To briefly summarize, cells were lysed in lysis buffer for 1 minute, followed by supplementation with 100% ethanol and vortexing for 10 seconds. The mixture was then transferred into a purification column at room temperature and spun down at 12,000 rpm for 1 minute at 4°C, and the supernatant was discarded. The column was washed with wash buffer A and B before being transferred into a new collection tube. Finally, 100 μl of nuclease-free water was added to the purification column, and it was spun down at 12,000 rpm for 1 minute to collect the flow-through. The cDNA was prepared using the QuantaBio qScript cDNA synthesis kit according to manufacturer’s instructions. To summarize, 0.1 μg of mRNA was combined with 1 μl of reverse transcriptase, 4 μl of reaction mix, and nuclease-free water was added to reach a final volume of 20 μl. The PCR reaction was run on a BioRad iQ5 thermocycler using iQ SYBR Green Supermix according to manufacturer’s instructions. Gene expression was calculated relative to 18S rRNA.

### Primers used for APOL7C expression

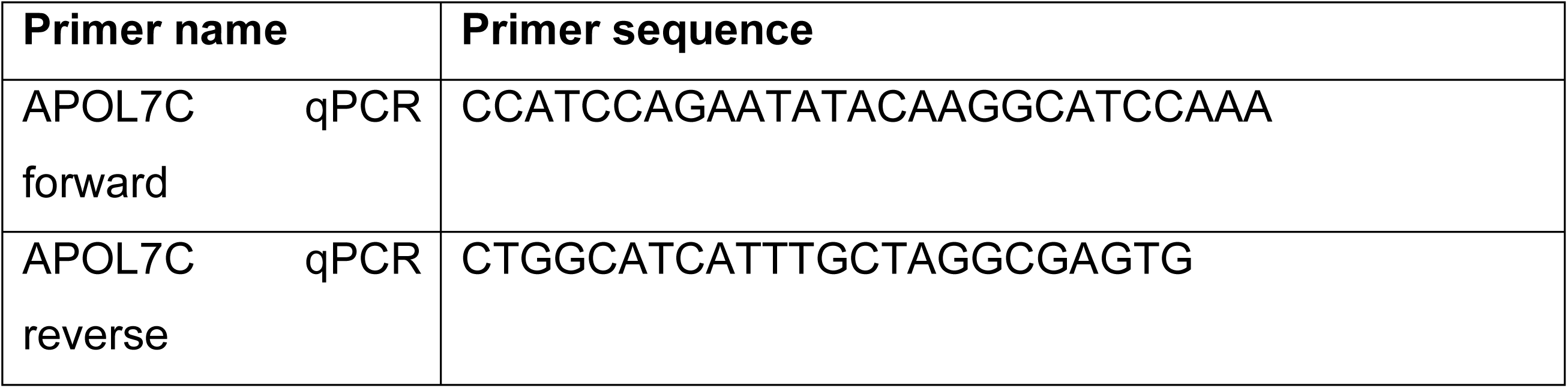

*Microscopy*. All imaging experiments were performed on a Leica SP8 DMi8 confocal microscope with LAS X MicroLab software. The microscope is equipped with an HC PL APO 63×/1.40 Oil CS2 objective, UV 405 nm laser, blue 488 nm laser, green 552 nm laser, red 638 nm laser, and PMT and HyD detectors. Cells were imaged on Ibidi 8-well μ-dishes in PBS (fixed or live cells) or on 12 mm glass coverslips mounted onto glass slides using ProLong Diamond mounting medium (fixed cells). All samples were imaged using Type F Immersion liquid at room temperature. Images were analyzed in Fiji (ImageJ2, version 2.9.0/1.53t, build a33148d777).

L. major *parasite maintenance.* The *L. major* (MHOM/IL/80)-red fluorescent protein (RFP)-expressing parasite was provided by Dr. Nathan C. Peters (University of Calgary). Parasites were grown at 26°C in medium 199 (M199) supplemented with 20% heat-inactivated FBS, 100 U/mL penicillin, 100 μg/mL streptomycin, 2 mM L-glutamine, 40 mM HEPES, 0.1 mM adenine (in 50 mM HEPES), 5 mg/mL hemin (in 50% triethanolamine), and 1 mg/mL 6-biotin. *L. major*-RFP^+^ cultures were further supplemented with geneticin antibiotic to maintain the plasmid.

In vitro L. major *infections and imaging*. FLT3L cDCs, MuTuDCs, MuTuDC.APOL7C::mCherry, RawKb, RawKb.APOL7C::GFP or RawKb.APOL7C::mCherry cells were plated onto Ibidi 8-well μ-dishes and left to adhere overnight. In the case of FLT3L cultures the cells were plated onto 12 mm glass coverslips that were coated with anti-MHC II. Just prior to infection, metacyclic *L. major* promastigotes were isolated from stationary cultures (4-5 days old) using a Ficoll gradient. The cells were then challenged with the metacyclic promastigotes incubated at 5% CO_2_ at 34°C for the time indicated and immediately fixed with 4% PFA in PBS. PFA was then quenched with 50 mM NH_4_Cl in PBS for 5 minutes followed by 3 washes with PBS. In some experiments, outside parasites were labeled with an anti-LPG antibody. Next, the cells were permeabilized with 0.4% Triton X 100 in PBS for 10 minutes followed by 3 washes with PBS. Cells were blocked for 30 minutes with 2% low fat milk in PBS and then incubated with the primary antibodies for 1-2 hours are room temperature. Cells were then washed 3X with 2% low fat milk in PBS. Secondary antibodies were then added for 1 hour, followed by 3 washes. Cells were then left in PBS for imaging (μ-dishes) or were mounted onto glass slides using ProLong Diamond mounting medium. Images were then acquired on a Leica SP8 confocal microscope.

*Luminol assay.* RawKb.APOL7C::mCherry cells were plated at a density of 1 X 10^5^ cells per well in a white 96-well clear bottom plates and incubated to adhere overnight. Cells were then washed 3X and bathed in PBS containing 10mM glucose, 50 μM luminol and 8U/mL horseradish peroxidase (HRP). Cells were then immediately infected with *L. major* at the indicated MOIs. The production of reactive oxygen species was monitored by analyzing chemiluminescence every 2 minutes for a total of 120 minutes at 34°C.

*Nitroblue tetrazolium assay.* RawKb.APOL7C::mCherry cells were plated onto Ibidi 8-well μ-dishes to adhere overnight. The next day, cells were washed 3X with serum free RPMI and infected with *L. major* (MOI 25) for 4 hours. NBT at 10 μg/mL was added to the medium after infection and cells incubated for 15 minutes before immediate brightfield imaging.

In vitro L. major *infection and IFN-γ detection.* FLT3L-cDCs were seeded at a density of 1 X 10^5^ cells per well in a 96-well U bottom plate. Metacyclic promastigotes of *L. major*-OVA were then added at the indicated parasite:cell ratio for 6 hours at 5% CO_2_ at 34°C. OT-I CD8+ T cells were added at a density of 2.5 X 10^4^ cells per well in RPMI supplemented with 10% HI-BGS, penicillin, streptomycin and β-mercaptoethanol. After 24 hours, the plates were centrifuged and the supernatant was collected and used for an IFN-γ ELISA as per instruction manual.

In vitro L. major *infection and IL-12 detection.* FLT3L-cDCs were seeded at a density of 1 X 10^5^ cells per well in a 96-well U bottom plate. Metacyclic promastigotes of *L. major* were then added at the indicated parasite:cell ratio overnight incubated at 5% CO_2_ at 34°C. After 24 hours, the plates were centrifuged and the supernatant was collected and used for an IL-12 ELISA as per instruction manual.

*Loading lysosomes with fluorescent.* RawKb.APOL7C::mCherry cells were plated onto Ibidi 8-well μ-dishes and left to adhere for 1 hour. Cells were then incubated with 10 kDa dextran conjugated to AF647 (125 μg/mL) to allow for the internalization of fluorescent dextran. After 4 hours of internalization, excess fluorescent dextran was washed off with RPMI supplemented with 10% HI-BGS, penicillin, and streptomycin. Cells were then treated with 1 μg/mL doxycycline overnight. The following day, cells were washed 3X with serum free RPMI and infected with *L. major* (MOI 25) for 3 hours. Cells were then washed 3X with serum free RPMI and immediately imaged live on a Leica SP8 confocal microscope described above.

*Lucifer Yellow (LY) and poly(dA:dT) leakage assays.* RawKb.APOL7C::mCherry or RawKb.APOL7C::GFP cells were treated with 1 μg/mL doxycycline and plated onto Ibidi 8-well μ-dishes to adhere overnight. The next day, cells were washed 3X with serum free RPMI and infected with *L. major* (MOI 25) and LY (150 μg/mL) for 6 hours, or incubated with non-porous silica beads with poly(dA:dT)-Rhodamine (30 μg/mL) for 5 hours. Cells were then washed 3X with serum free RPMI and immediately imaged live on a Leica SP8 confocal microscope described above.

*Coating coverslips with anti-MHC II.* Glass coverslips (12 mm) were covered in a 0.1 % solution of poly-L-lysine for 30 minutes at room temperature. Coverslips were then washed 3X with PBS and incubated in 2.5 % glutaraldehyde for 15 minutes at room temperature. Coverslips were then washed 3X with PBS. A solution of 1 μg/mL of anti-MHC II in PBS was then overlayed on the coverslips for 30 minutes at room temperature. Coverslips were then washed 3 more times with PBS and then incubated overnight in 0.2 M glycine in PBS at 4°C. On the day of the experiment, coverslips were washed an additional 3 times with PBS prior to attaching cells.

*Flow cytometry.* Single cell suspensions were aquired for infected ears and cervical draininf lymph nodes. Ventral and dorsal ear sheets were separated and digested with 16μg/mL of Liberase (Roche Diagnostic) in 0.5 mL DMEM for 90-120 minutes at 37°C. Digested ear tissues were then homogenized for 40’ at 1714 rounds per run (rpr) using a gentleMACS Octo dissociator instrument (Miltenyi Biotec,Inc) with 6 ml DMEM media containing 0.05% DNase I and filtered using a 50 μm-pore-size cell strainer. Ear draining lymph nodes (dLNs) were removed and homogenized with a 1mL syringe plunger on a 70μm cell strainer. After tissue processing cells were washed and labeled with Live/Dead fixable viability stain and anti-Fc III/II (CD16/32) receptor Ab (2.4G2) for 20 min at 4°C. This step was followed by a surface staining with different combinations of antibodies for 20 min at 4°C in the dark. For intracellular staining, cells were fixed with BD Cytofix/Cytoperm (BD Biosciences) and stained for 45 min at 4°C when staining for cytokines. For the analysis of intracellular protein phosphorylation cells were immediately fixed (BD Cytofix Buffer) at 37°C for 10m, followed by a permeabilization (BD Phosflow Perm Buffer II) step at 4°C for 30m and then stained with both surface and phosphor-specific markers. Cells were washed between the steps of fixation-permeabilization and final staining. All data acquisition was performed using a 5-laser Northen Data were collected from individual ears using a 5-laser Cytek Aurora (Cytek Biosciences) spectral flow cytometer and analyzed using FlowJo software (TreeStar). The follow anti-mouse antibodies were used: CD11b (M1/70), CD45 (30-F11); Ly6G (1A8); Ly6C (HK1.4); CD64 (X54-5/7.1); CCR2 (SA203611); CD11c (HL3); MHCII (M5/114.15.2); CD90.2 (53-2.1); TCR-b (H57-597); CD4 (RM4-5); CD8 (53-6.7); IFN-γ (XMG1.2); SiglecF (1RNM44N); XCR1 (ZET); CCR7 (4B12); CD24 (M1/69); CD49b (HMa2); CD200R3 (Ba13); CD172a (P84); IL-12p40/p70 (C15.6); T-bet (eBIO 4B10), GATA3 (L50-823); RORdT (B2D); STAT4p (pY693); KI-67 (So1A15). All antibodies were obtained from Thermo Fisher, BD Biosciences or Biolegend.

*Correlative Focused Ion Beam Scanning Electron Microscopy (FIB-SEM).* RawKb.cGFP.APOL7C::mCherry cells were seeded into a gridded glass ibidi μdish at 20% confluency. The culture medium was supplemented with doxycycline to a final concentration of 1 μg/ml and incubated overnight at 37°C. The next day, cells were thoroughly washed and incubated with *L. major* (MOI 25) for 4 hours before imaging at 37°C, 5% CO_2_ on Leica SP8 confocal microscope equipped with a HC PL APO 63×/1.40 Oil CS2. Following confocal imaging, the ibidi dish was washed twice with PBS to remove the culture medium and immediately fixed with 2% PFA + 2.5% glutaraldehyde in 0.2 M cacodylate buffer for 1 hour. The cells were then washed three times with 0.15 M cacodylate buffer and stained with 2% OsO_4_ in 1.5% potassium ferrocyanide-containing cacodylate buffer for 90 minutes. The ibidi dish was washed three times with 0.15 M cacodylate buffer and dehydrated with a series of ethanol concentrations (50%, 70%, 90%, 95%, and 100%) and pure acetone for 15 minutes each. The Durcupan resin was prepared by combining part A (11.4g), part B (10g), part C (0.3g), and part D (0.1g). For better infiltration, the resulting resin was mixed with acetone in ratios of 3:1, 1:1, and 1:3, and added sequentially to the ibidi dish after acetone dehydration for infiltration, with each step taking 30 minutes and a final overnight step at 4°C with 100% resin. On the following day, the overnight resin was replaced with fresh resin and polymerized in a 60°C oven for 48 hours.

The serial sectioning and imaging was performed on a dual-beam FIB-SEM Helios 5 CX (Thermo Fisher Scientific Inc.) equipped with a gallium liquid metal ion source. The resin blocks were mounted on an SEM stub using a conductive epoxy resin glue and with silver resin paste and then sputter-coated with carbon (Leica EM ACE600) to enhace sample conductivity before being loaded onto a multi-purpose holder in the SEM sample chamber. An overview low resolution back scattered electron (BSE) image of the entire block was acquired with the Everhart-Thornley detector to identify cells of interest (COI) by comparing with the confocal image. Once the COI was identified, the sample was positioned at the eucentric height and was prepared for the serial-sectioning and imaging (FIB-BSE) workflow. Briefly, a ∼1–2 µm thick protective layer of platinum was deposited on the top of the COI using a gas injection system (GIS) and trenches (∼60 µm (X) × 30 µm (Y) × 15 µm (Z)) were dug in front and along each side (∼ 10 µm wide) of the COI using the focused ion beam (FIB) to expose its block face. Once prepared, the COI was sequentially milled using the FIB and imaged using the SEM and BSE signals via an automated serial sectioning and imaging workflow (Auto Slice & View 4.2 (ASV) software package). SEM and BSE images were acquired using 2kV accelerating voltage, x mm working distance, and at a beam current of 0.69 nA with the Elstar in-lens (TLD) and the Elstar in-column (ICD) detectors, respectively, and typical dwell times of 3-5 μs. This allowed us to obtain high-resolution three-dimensional (3D) images (6-8 x 6-8 x 10 nm^3^ voxels) of the COI. The 3D images were then aligned and segmented manually using the TrakEM2 plugin in ImageJ (version 2.9.0/1.53t, build a33148d777) and registered and reconstructed in Imaris 10.0.0 (Oxford Instruments).

*In vivo infections.* C57BL6 Wt and APOL7C^-/-^ mice infected with *Leishmania major* Friedlin (FV1)-RFP parasites. Mice either received 10^4^, for the long-term course of infection (7 and 18 weeks) or 2x10^5^, for the short-term experiments looking at CD4 T cell priming response (48h), *L. major* metacyclic promastigotes i.d in the ears in a volume of 10μl. For the long-term experiments ear lesion size was weekly measured with the use of a caliper. Parasite numbers were determined by limiting dilution analysis (LDA), where two-fold serial dilutions in a 96-well flat bottom polystyrene microtiter plates were performed using M199 complete medium. After 10 days of incubation at 26°C, numbers of parasites per sample was determined from the highest dilution well at which parasites were present. Ethics approval for this study was obtained from the Animal Care Committee (ACC) at the University of Calgary (Protocol Number: AC23-0019). Mice were anesthetized using Ketamine-Xylazine prior to ear infections.

*Single-cell RNA-Seq bioinformatics.* Filtered feature-barcode HDF5 matrices from the dataset downloaded from the NCBI GEO (GSE181720) were imported into the R package Seurat v.4.3.0 (for normalization, scaling, integration, Louvain clustering, dimensionality reduction, differential expression analysis, and visualization^56,57^. Briefly, cells with fewer than 1000 features, greater than 2,500 features, and greater than 5% of mitochondrial reads were excluded from subsequent analysis. Cell identity was annotated based on previously published markers^23,24^.

*Statistical analysis.* GraphPad Prism (version 10) software was used to perform all statistical analyses. Statistical significance comparing two samples was determined using one way analysis of variance (ANOVA) or unpaired, two-tailed Student’s *t* test with or without Welch’s correction, Kolmogorov-Smirnov test, or unpaired parametric Mann-Whitney tests as indicated in the figure legends. Two-way ANOVA was used for experiments which had two independent variables. Data are displayed as mean ± standard deviation or ± standard error of the mean as indicated in the figure legend. Results were considered statistically significant at *P* < 0.05.

---

**S1 Fig.** Gating strategy for *L. major* infected cDCs. Related to. Fig 1**. (A)** Single cell suspensions from the skin of *L. major* infected ears of C57BL/6J mice assessed for cDC populations by flow cytometry. cDC1s are considered to be CD11B^-^, CD11C^+^, MHCII^+^, XCR1^+^, monocyte-derived dendritic cells are CD11B^+^, LY6G^-^, LY6C^-/Lo-Hi^, CD64^+^, CCR2^+^, MHCII^+^, and Het-DCs are considered to be LY6C^-^, LY6G^-^, CCR2^+/-^, CD64^-^, CD11C^+^, MHCII^+^. **(B)** Gating strategy for infected *L. major* RFP+ cells. Shown are representative plots from 2 independent experiments of mice infected with *L. major* for 7 weeks.

**S2 Fig.** APOL7C positive PVs harbour breaks in L. major membranes. Related to. Fig 3**. (A)** RawKb.cGFP.APOL7C::mCherry cells were plated and treated with doxycycline overnight and challenged with *L. major* at an MOI of 25 for 4 hours. A cell with an APOL7C-positive PV was then processed for FIB-SEM. A single FIB-SEM slice of a cell with an APOL7C-positive PV is shown and annotated for PV membrane and *L. major* parasite membrane.

**S3 Fig.** APOL7C-deficient cells exhibit higher parasite burden. Related to. Fig 3**. (A-B)** FLT3L cDCs derived from *Apol7c^+/+^* and *Apol7c^-/-^* mice were plated and challenged with RFP-*L. major* at an MOI of 50 for 6 hours. Cells were then fixed and stained with Galectin 3 to visualize the cytosol and a nuclear stain. **(B)** The number of internal RFP-*L. major* parasites per cell was quantified. Data is plotted as mean ± SEM of three independent experiments (n=3), significance determined using student’s *t* test. **(C-E)** FLT3L cDCs derived from *Apol7c^+/+^* and *Apol7c^-/-^* mice were plated and challenged with RFP-*L. major* at an MOI of 50 for 6 hours. **(C)** Total CD45+B220-CD11c+MHCII+ cDCs, **(D-E)** CD45+B220-CD11c+MHCII+XCR1+ cDCs positive for RFP- *L. major* was quantified. Data is plotted as mean ± SEM of three independent experiments (n=3), significance determined using student’s *t* test. DIC; differential interference contrast, n.s. = no significance; \**P* £ 0.05; \*\*\**P* £ 0.001.

**S4 Fig.** Parasite numbers at the site of infection and draining lymph node and analysis of TH1 cells at the draining lymph node. Gating strategy for TH1 and TH2 T cell mediated immunity. Related to Fig 4. (A-B) RawKb.APOL7C::GFP cells were left untreated or treated with LPS (500 ng/mL) for 1 hour. Cells were then fixed and stained for NFΚB along with a nuclear stain before imaging with confocal microscopy. (G) The nuclear stain was used to mask and quantify the mean fluorescence intensity (MFI) of NFKB in the nucleus. Data is plotted as mean ± SEM of three independent experiments (n=3), significance determined using student’s *t* test. (C-D) RFP-*L. major* parasite load of intradermally inoculated *Apol7c^+/+^* and *Apol7c^-/-^* mice in the (C) skin and (D) draining lymph node. Plotted as mean ± SEM. Each dot represents a single mouse. Welch’s *t* test between *Apol7c^+/+^* and *Apol7c^-/-^* mice was performed to determine significance. (E-F) Gating strategy for stained single cell populations from the draining lymph node of RFP-*L. major* infected mice. Shown are representative plots from 2 independent experiments. **(E)** IL-12 producing cDC1s are considered to be CD45+, CD11b-, MHC-II+, CD90.2-, CD172a-, XCR1+, and IL12+. **(F)** CD4+ T cells are considered to be CD45+, CD11b-, CD90.2+, MHC-II-, CD8-, and CD4+. **(G-H)** Gating strategy of naive and infected CD4+ T cells that are **(G)** CD45+ and CD90.2+, stained for T bet, GATA3 and RORγT and **(H)** CD45+, CD90.2+, and Tbet+ cells stained for STAT4p, IFNγ and Ki67. Shown are representative plots from 2 independent experiments. TL = transmitted light; n.s. = no significance; \*\*\*\**P* £ 0.0001.

S1 Movie. APOL7C::mCherry is recruited to *L. major* containing PVs. Related to Fig 2**. (Left)** RawKb.APOL7C::mCherry cells were transfected with F-tractin::GFP and treated with doxycycline overnight. Cells were then washed and infected with *L. major* parasites at an MOI of 20 for 4 hours before live cell confocal imaging. **(Right)** RawKb.cGFP.APOL7C::mCherry cells were plated and treated with doxycycline overnight. Cells were then washed and infected with *L. major* parasites at an MOI of 20 for 4 hours before live cell confocal imaging.

**S2 Movie. APOL7C-negative PVs exhibit continuous membranes. Related to** Fig 3**. (Left)** Live cell confocal imaging and **(Right)** Segmented FIBSEM of a RawKb.cGFP.APOL7C::mCherry cell plated and treated with doxycycline overnight. Cells were then washed and infected with *L. major* parasites at an MOI of 20 for 4 hours before live cell confocal imaging and subsequent FIB SEM sample processing.

S3 Movie. APOL7C-positive PVs exhibit discontinuous membranes. Related to Fig 3**. (Left)** Live cell confocal imaging and **(Right)** Segmented FIBSEM of a RawKb.cGFP.APOL7C::mCherry cell plated and treated with doxycycline overnight. Cells were then washed and infected with *L. major* parasites at an MOI of 20 for 4 hours before live cell confocal imaging and subsequent FIB SEM sample processing.

## Notes

### Competing Interest Statement

The authors have declared no competing interest.

